# A novel anti-climbing barrier prevents black soldier fly larval escape from rearing containers

**DOI:** 10.64898/2026.04.28.721252

**Authors:** Sunjae Jang, Masami Shimoda

## Abstract

The mass-rearing of black soldier fly (*Hermetia illucens*) larvae (BSFL) is a promising solution for converting organic waste into high-quality insect protein, but preventing larval escape from open-top rearing containers remains a major management challenge. Conventional escape-control methods are often unreliable or impractical. To address this, we developed and evaluated a novel physical barrier, the anti-climbing tape, featuring regularly arranged macroscale protrusions designed to disrupt larval locomotion on vertical surfaces. We conducted a series of experiments to examine the design parameters of the anti-climbing tapes, including the gap size between protrusions and the number of protrusion rows. Our results demonstrate that the anti-climbing tape prevents escape via a dual mechanism: (1) physical obstruction, in which gaps narrower than the larval body width block larvae from passing through, and (2) adhesion reduction, in which the elevated protrusion array decreases the effective contact area for wet adhesion while increasing gravitational torque acting on the larval body. The effectiveness of these mechanisms was dependent on larval size. A design featuring 0.5-mm gaps and a 15-row protrusion array completely prevented the escape of later-instar larvae (>10 mm) in a 20-day large-scale trial, whereas the method was less effective for smaller larvae. In conclusion, the anti-climbing tape provides a robust and chemical-free approach to BSFL escape in mass rearing. To ensure reliable performance, its design parameters, both gap size and array width must be optimised to suppress the mechanical and adhesive components of larval climbing according to the target larval size.

**Conflict of interest:** S. Jang and M. Shimoda are inventors on a Japanese patent application (No. 2022-172252, filed November 27, 2022) related to the method described in this manuscript.

**Funding:** This study was supported by Korea-Japan Joint Government Scholarship Program for the Students in Science and Engineering Departments, the Korean Scholarship Foundation, and the University of Tokyo Foundation’s Support Fund for International Students.

## 1 Introduction

Insects are gaining international attention in recent years for two beneficial uses: (1) the source of human food or livestock feed (van Huis, 2020) and (2) the agent for treating organic wastes (Čičková *et al*., 2015; Mannaa *et al*., 2024). Among them, the black soldier fly (BSF), *Hermetia illucens* (Diptera: Stratiomyidae) larvae (BSFL) is emerging as a promising solution — BSFL devour a wide range of organic matters, including meat, vegetables, and even fecal wastes (Banks *et al*., 2014; Michishita *et al*., 2023), and produce high-quality protein-rich biomass (Barragan-Fonseca *et al*., 2017). For the successful incorporation of the abilities of BSFL into the resource circulation network of our society, stable mass-rearing of BSFL in a controlled environment is crucial (Cammack and Tomberlin, 2023). By nature, BSFL spend their lifetime burrowed inside feed. The depth of feeding substrate that can be penetrated and processed by larvae is limited to approximately 5-10 cm (Biasato *et al*., 2024; Čičková *et al*., 2015), as BSFL need to expose their spiracles in the air to breathe. Due to such limitations, current mass-rearing of BSFL on an industrial scale mainly adopts shallow containers with a large surface area (Liu *et al*., 2022; Yang and Tomberlin, 2020). This type of rearing setup is conventionally managed without covering lids for effective ventilation to prevent over-humidification within containers (Dortmans *et al*., 2021), and for management convenience, which is inherently vulnerable to larval escape.

Although no study has extensively investigated the cause of BSFL escape from artificial environment to our knowledge, possible factors can be categorised into three groups: (1) unfavorable environment — *e.g*., excessive heat (Fuhrmann *et al*., 2025), anaerobic condition (Pham, 2022), or intense light (Giannetti *et al*., 2022); (2) abrupt environmental change — *e.g*., placing larvae into a fresh batch of feed (Jang, personal observation); and (3) innate programmed-migration at prepupal stage in search of dry environment (Sheppard *et al*., 1994). Nevertheless, the escape behavior of BSFL remains somewhat enigmatic, as unusually small individuals, possibly due to malnutrition, are often observed outside the feed even under conditions considered optimal. The larval escape may lead to several non-negligible problems in rearing management, such as dispersion of larvae, detriment of environmental hygiene, and overall increment in the burden of management for workers. Moreover, if escaped larvae succeed in metamorphosis to adults out of human attention, gaining increased mobility, this will raise the risk of accidental leakage of genetic resources into the wild ecosystem (Tepper *et al*., 2024).

Like most other dipteran larvae, BSFL possess no appendages or structures for locomotion (*i.e*., legs and prolegs; Barros *et al*., 2019). However, in the presence of viscous fluid (*e.g*., water), via wet adhesion, larvae can adhere and crawl on smooth surfaces, including glass and plastics, positioned vertically or even upside down. There are no reports that BSFL themselves secrete fluid for adhesion, as is the case for legged adults of various insect taxa (Dirks and Federle, 2011), gastropods (Smith, 2010) or tree frogs (Langowski *et al*., 2018). As BSFL instantly become unable to adhere to such surfaces when the fluid is removed, it is obvious that larvae require fluid of external origin for climbing and further escape. Based on such empirical facts, conventional approaches to prevent BSFL escape have centered on controlling fluid or moisture in the rearing environments. Examples include placing dry substances to soak up moisture from larval body surface (Dortmans *et al*., 2021) and lowering the moisture content of the feeding substrate itself (Vandeweyer and Van Der Borght, 2022). However, these methods have respective limitations: powder loses its effect as it becomes wet and requires replacement; controlling the feed moisture has been reported to directly affect larval feeding and growth (Bekker *et al*., 2021; Dzepe *et al*., 2020). In addition, the use of chemicals such as repellents or insecticides — a conventional method for controlling insect migration — is not a viable option, considering the potential risk of their bioaccumulation in the final larval products (Meijer *et al*., 2021).

To address these limitations, we developed a novel method that prevents BSFL climbing and subsequent escape from rearing containers using custom-designed, tape-like structures (Figure 1). These structures, hereafter referred to as ‘anti-climbing tapes’, bear regularly arranged macroscale protrusions that disrupt larval locomotion on vertical surfaces. The anti-climbing tapes are readily applicable to conventional BSFL rearing containers without structural modification and effectively suppress larval escape even in high-moisture environments.

**FIGURE 1.**
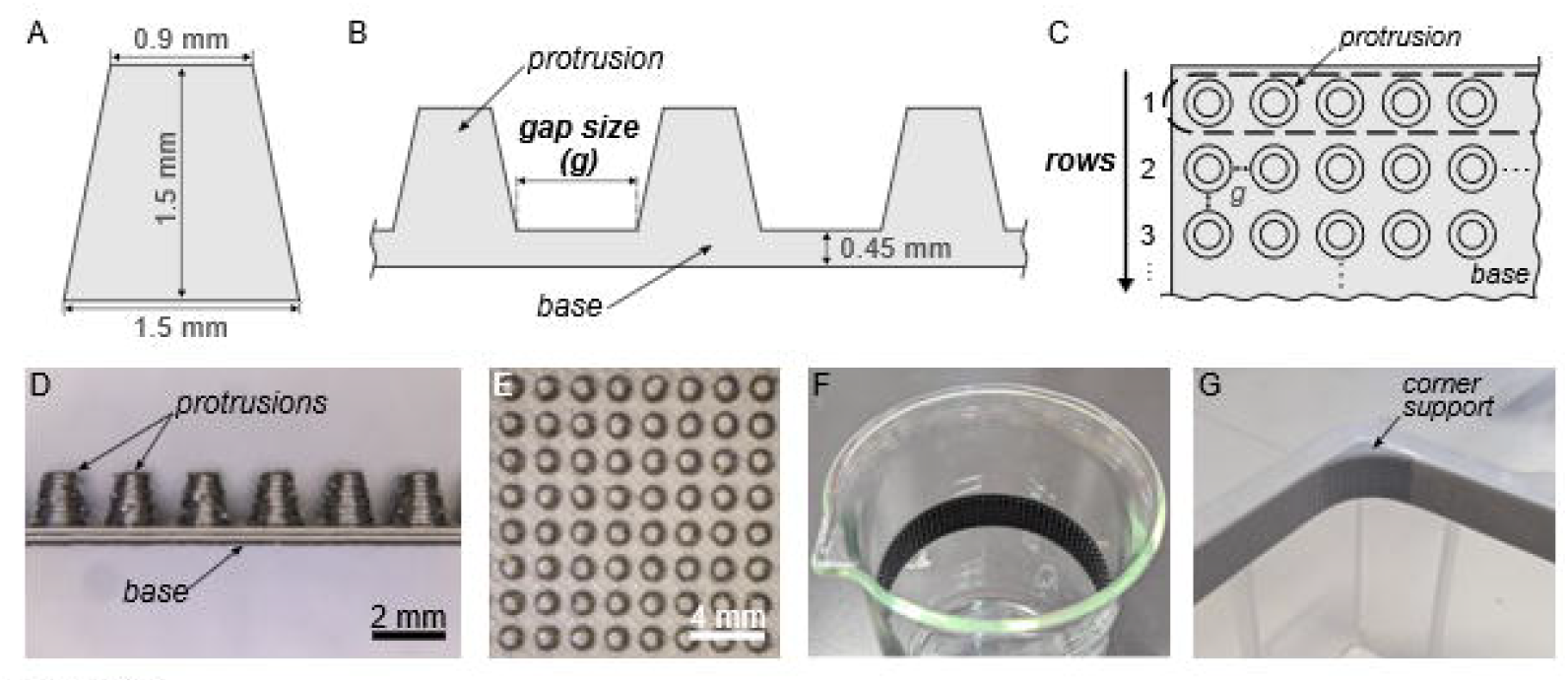
Diagrams and photographs of anti-climbing tapes. (A) Side view of a protrusion. (B) Definition of gap size between protrusions. (C) Definition of a row in a protrusion array. (D) Side and (E) top vie of a 3D-printed anti-climbing tape with 0.5-mm gap size. Examples of the tape application on (F) a cylindrical glass beaker and (G) rectangular polypropylene container. Wedge-like supporting structures were used for the comers of rectangular containers.

## 2 Materials and methods

### Insect rearing

For adhesive measurements, the Tsukuba (TK) strain of BSF was used. The TK strain was established from wild flies collected at Tsukuba City, Ibaraki Prefecture, Japan in 2013 (Nakamura *et al*., 2016). Adult flies were reared following the method described by Nakamura *et al*. (2016), in a greenhouse at 35 ± 10°C, 55 ± 25% relative humidity (RH). Collected eggs were transferred to a plastic cup and incubated at 28 ± 2°C, 62.5 ± 12.5% RH, and 16L:8D photoperiod. Hatched larvae were fed with a diet containing soy pulp, rice bran, and potato starch (DW), and kept at 15°C. Larvae that reached approximately 20 mm in length and 100 mg in mass were used for the measurement. Based on their body mass and head capsule morphology, these larvae were considered to be at fifth instar (Fabian *et al*., 2025; Kim *et al*., 2010)

For escape-prevention experiments, in addition to the TK strain, the Kagoshima (KG) strain was used. The KG strain was established from wild flies collected at Kagoshima City, Kagoshima Prefecture, Japan in 2020 (Ohki *et al*., 2025). For both strains, adult flies were reared in the same manner described above. Collected eggs were transferred to a plastic cup and incubated at 25 ± 2°C, 60 ± 10% RH, and 12L:12D photoperiod. The eggs were checked for hatched larvae at 24-hour intervals. Larval age, in day-old (DO), was counted from the day hatching was confirmed (day 0). Larvae were fed with an artificial diet containing glucose, molasses, dry yeast, cornmeal, and agar (Liu *et al*., 2024), and incubated under the same conditions as above. The larvae of the TK strain (10 and 20 DO) were used for Experiment 1 to 3, while the larvae of the KG strain (10 DO) were used for Experiment 4. Larval body length was approximately 10 mm for 10-DO and 20 mm for 20-DO larvae. Based on the body length, larval groups were considered to consist of fourth to fifth instars for 10-DO and fifth to sixth instars for 20-DO (Barros *et al*., 2019; Fabian *et al*., 2025).

### Adhesive force measurements

Using a custom-made device incorporating a load cell (Supplementary Figure S1, 2), we measured the force generated by wet adhesion between larvae and substrate samples. Before measurement, each larva was freeze-killed and fixed dorsally on a plate for measurement with cyanoacrylate glue (Supplementary Figure S1A). To apply water for adhesion on the larvae, the center of the ventral side of the larvae was pressed vertically onto a 5 µL droplet of DW on a borosilicate slide glass (S1126, Matsunami Glass Ind., Ltd., Osaka, Japan) under a load of 1.0 ± 0.2 mN and vertically pulled off at the speed of 1 mm/s, three times. To measure the force, each larva with water on it was pressed onto a substrate for six seconds and pulled off at the same load and speed mentioned above, respectively. From the peak value of the forces observed right after the removal of 1 mN load, we subtracted a baseline value (mean of thirty consecutive values recorded one second after the peak value; Supplementary Figure S1C). The resulting value was regarded as ‘adhesive force’ and standardised by dividing by larval weight (mass × gravitational acceleration).

For substrate samples, commercially available materials and 3D-printed anti-climbing tapes were used for force measurements.

First, five commercially available materials were prepared: (1) borosilicate slide glass as control, (2) polytetrafluoroethylene (PTFE) tape (ASF-110FR, Chukoh Chemical Industries, Ltd., Tokyo, Japan) for its smooth and hydrophobic surface (Dhanumalayan and Joshi, 2018), (3) sandpaper (150 grit; DCCS-150, Sankyo Rikagaku Co., Ltd., Okegawa, Japan) for its granular surface with microscale protrusions (20-200 μm in height; Crawford *et al*., 2016; Stief *et al*., 2020), and (4,5) hook-and-loop fastener tapes (hook and loop components were tested separately; G511N (hook) and G521N (loop), Kuraray Fastening Co., Ltd., Tokyo, Japan) for their macroscale protrusions (2–3 mm in height). Materials except control were fixed on a slide glass, separate from measurement samples, using double-sided adhesive tape. Using 15 larvae, one measurement was taken per individual on each material, in a randomised order between materials.

Second, the anti-climbing tapes were fabricated by fused filament fabrication (FFF) 3D printing with polylactic acid (PLA) filament (Pro3 Plus and PLA filament, Raise 3D Technologies Inc., Irvine, CA, USA) at 0.15 mm layer height (Figure 1). Basically, each tape consisted of a flat rectangular base (0.45 mm thick) with truncated cone-shaped protrusions (1.5 mm high, 0.9/1.5 mm top/bottom diameter; Figure 1A). Two types of tape were prepared: (1) base-only tapes without protrusions and (2) tapes with protrusions arranged in a rectangular grid, spaced at 0.5 mm apart (0.5 mm gap). Each tape was fixed on a slide glass with double-sided adhesive tape. Using four larvae, one to two measurements were taken per individual on each tape, in a randomised order between tapes.

### Escape-prevention experiments

The escape-prevention effect of the commercially available materials and custom-designed anti-climbing tapes on BSFL was evaluated in a series of experiments.

Experiment 1: Using four commercially available materials, PTFE, sandpaper, hook, and loop, larval escape-prevention from rearing containers was attempted. On the inner surface of the 500 mL cylindrical borosilicate glass beaker (inner dimensions: diameter 85 mm × height 120 mm; S75-1001-25, AGC Techno Glass Co., Ltd., Tokyo, Japan), each material was applied as a 20-mm-wide, continuous circumferential band. The lower edges of the bands were positioned 55 mm above the bottom. Tape ends were seamlessly joined, and double-sided adhesive tape was used for attachment. Untreated containers served as control. For each beaker, 20 larvae were placed with 10 mL of artificial diet diluted with distilled water (DW) as substrate (diet:DW = 1:3 in mass). The beakers were placed inside rectangular containers containing dried rice husk or sawdust to trap escaped larvae. After 24 and 48 hours, 10 mL of DW was added to each beaker to provide moisture, since the desiccation of substrate hindered larval climbing in preliminary trials. After 72 hours, the number of larvae remaining in each beaker was counted, and the larval escape rate was calculated as:

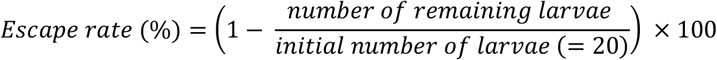

Trials were conducted separately for two age groups (10-DO and 20-DO), with five replicates per treatment and age group. Each beaker contained larvae of only one age. Environmental conditions were controlled at 25 ± 2°C, 60 ± 10% RH, and 12L:12D photoperiod.

Experiment 2: Using 3D-printed anti-climbing tapes with different protrusion gap sizes, larval escape-prevention from rearing containers was attempted. On a 20 mm-width base, protrusions were arranged in square arrays with the gap size of 0.5, 1.5, or 2.5 mm (Figure 1B), and the arrays were aligned symmetrically with the base midline. To maintain consistent spacing of protrusions at which the tape ends were joined, half-gap offsets were applied at both ends. The number of protrusion rows (Figure 1C) varied among experiments. Base-only tapes were also prepared. Design parameters of the anti-climbing tapes are summarised in Table 1. As in Experiment 1, each tape was applied onto separate glass beakers. Containers untreated and treated with base-only tape served as control. Using the above containers, the experiment was conducted in the same manner and environmental condition as described in Experiment 1 section.

**TABLE 1.** Design parameters of anti-climbing tapes.

|  | Base width (mm) | Gap size (mm) | Row number |
| --- | --- | --- | --- |
| Experiment 2 | 20 | - | 0<sup>1</sup> |
|  |  | 0.5 | 10 |
|  |  | 1.5 | 6 |
|  |  | 2.5 | 5 |
| Experiment 3 | 20 | - | 0<sup>1</sup> |
|  |  | 0.5 | 1 |
|  |  |  | 4 |
|  |  |  | 7 |
|  |  |  | 10 |
| Experiment 4 | 30 | 0.5 | 15 |

<sup>1</sup>Tapes without protrusions (base only).

Experiment 3: Using anti-climbing tapes with different protrusion row numbers, larval escape-prevention was attempted. Tapes (20 mm wide, 0.5 mm gap) with protrusion row numbers of 1, 4, 7, and 10 were prepared and applied to beakers. Design parameters of the tapes are summarised in Table 1. Containers treated with base-only tape served as controls. Using the above containers, the experiment was conducted in the same manner and environmental condition as described in Experiment 1 and 2 sections.

Experiment 4: Under a mass-rearing setup corresponding to industrial scale, the escape-prevention performance of anti-climbing tape was tested. Tapes with a 30 mm-width base and protrusions arranged in 15 rows with 0.5-mm gaps were prepared and applied to rectangular polypropylene container (inner dimensions: width 325 mm × depth 443 mm × height 163 mm; Sanbox #23-3, Sanko Co., Ltd., Shiojiri, Japan). To ensure smooth curving of the tapes at the inner corner of the containers, 3D-printed wedge-like supports (30 mm wide curved surfaces; radius 38.44 mm) were installed beneath the tapes at all four corners. Gaps between supports and walls were sealed with foam clay. Untreated containers served as control. At the start of the experiment (day 0), 650 larvae (10 DO) were placed into each container filled with a mixture of 400 g breadcrumbs (Keiai Science, Tsukuba, Japan), 400 g rice bran (provided by a local rice shop, Tokyo, Japan), and 3,200 mL tap water as a substrate. Each container was placed in a larger container (inner dimensions: w428 mm × d648 mm × h163 mm; Sanbox #45-4, Sanko Co., Ltd., Shiojiri, Japan) containing dried sawdust. During 20 days of rearing, escaped larvae were collected daily from sawdust, and the feeding substrate was gently stirred with a silicone spatula. On day 7, 500 mL tap water was added to each container. The cumulative larval escape rate per container was calculated as:

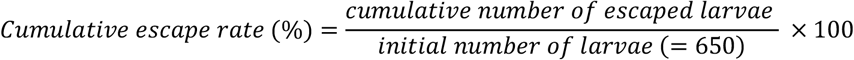

Three replicates were conducted per treatment. Environmental conditions were controlled at 28 ± 2°C, 62.5 ± 12.5% RH, and 16L:8D photoperiod, although the dark phase on day 6 was briefly disrupted by an 8-s light exposure one hour after its onset.

### Statistical analyses

Adhesive force measurements: Standardised adhesive force was analysed separately for commercially available materials and 3D-printed anti-climbing tapes. First, for commercially available materials, a generalised linear mixed effects model (GLMM) was applied, with material treatment and individual larva as fixed and random effects, respectively. Residuals were standardised. Overall differences among materials were assessed using a Type III Wald χ^2^ test, followed by Tukey’s post hoc comparisons based on general linear hypotheses. For 3D-printed anti-climbing tapes, Welch’s *t*-test was used. Normality of the data was confirmed by Shapiro-Wilk test. Although homoscedasticity was not rejected by Levene’s test (*P* = 0.065), Welch’s *t*-test was selected due to marginal variance inequality.

Escape-prevention experiments: Differences in larval escape rate between treatments were analysed using pairwise Wilcoxon rank-sum tests, conducted separately for each age group. *P*-values were derived from the normal approximation of the *W* statistics and adjusted using the Benjamini-Hochberg method.

All analyses were conducted in R (version 4.5.1; https://www.r-project.org/). No statistical analyses were performed for larval-escape data in Experiment 4.

## 3 Results

### Adhesive force measurements

In measurements using commercially available materials, we confirmed significant differences in standardised adhesive force among materials (Figure 2A; Wald χ^2^(4) = 48.481, *P* = 7.491 × 10^-10^; see Table 2 for *P*-values). The PTFE exhibited approximately 42% of the mean force of the glass control. In contrast, the sandpaper and hook-and-loop fasteners exhibited lower adhesive forces than both the glass control and the PTFE. Their adhesive forces corresponded to approximately 4–24% of the mean force of the glass control. No significant difference was detected among the sandpaper, hook, and loop.

**FIGURE 2.**
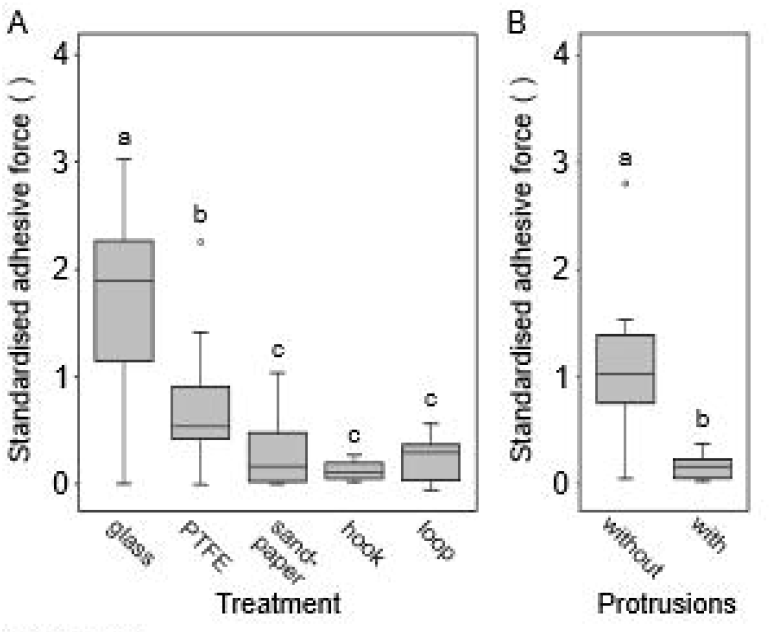
Adhesive force via wet adhesion between black soldier fly larvae (BSFL) and substrates. (A) Commercially available materials. (B) 3D-printed anti-climbing tapes. Data were presented as mean ± standard error (A: *n* = 15; B: *n* = 7). All force data were divided by larval body weight (body mass × gravitational acceleration), hence dimensionless. Different letters above bars indicate significant differences (*P* < 0.05) according to Tukey's multiple comparisons following a generalised linear mixed effects model (for A) or Welch’s t-test (for B).

**TABLE 2.** Summary of multiple comparisons among adhesive forces between BSFL and commercially available materials. *P*-values derived from Tukey’s multiple comparisons following a generalised linear mixed effects model are shown. **:*P*< 0.01. ***: *P* < 0.001.

|  | glass | PTFE | sandpaper | hook |
| --- | --- | --- | --- | --- |
| PTFE | 0.01** |  |  |  |
| sand-paper | < 0.001*** | 0.006** |  |  |
| hook | < 0.001*** | < 0.001*** | 0.391 |  |
| loop | < 0.001*** | 0.002** | 0.986 | 0.659 |

In measurements using 3D-printed anti-climbing tapes, we confirmed that macroscale protrusions regularly arranged on the surface significantly reduced the adhesive force to approximately 14% of the mean force measured on the flat surface without protrusions (Figure 2B; *t* = -3.064, *df* = 6.273, *P* = 0.021).

### Escape-prevention experiments

In Experiment 1, we confirmed a strong escape-prevention effect from containers treated with either segment of commercially available hook-and-loop fastener (Figure 3; see Table 3 for *P*-values).

**FIGURE 3.**
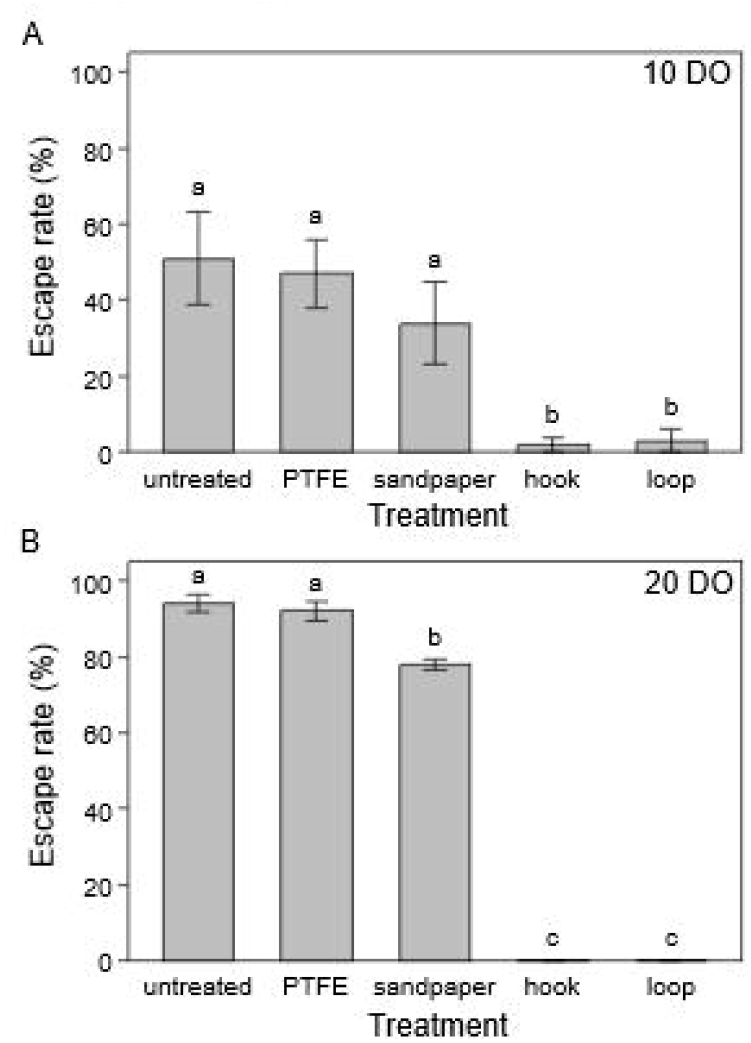
Escape rates of BSFL from containers treated with commercially available materials. (A) 10-day-old (DO) larvae. (B) 20-DO larvae Data were presented as mean ± standard error (*n* = 5). 20 individuals were used per replication. Different letters above bars indicate significant differences (*P* < 0.05) according to pairwise Wilcoxon rank sum test with Benjamini-Hochberg correction.

**TABLE 3.** Summary of multiple comparisons between larval escape rates from rearing containers treated with different commercially available materials. *P*-values derived from pairwise Wilcoxon rank sum test with Benjamini-Hochberg correction are shown. Values inside parentheses indicate *P*-values without correction. *: *P*< 0 05, NA: test not available (complete coincidence of values between groups)

|  | 10 DO |  |  |  |  | 20 DO |  |  |  |
| --- | --- | --- | --- | --- | --- | --- | --- | --- | --- |
|  | untreated | PTFE | sandpaper | hook |  | untreated | PTFE | sandpaper | hook |
| PTFE | 1.000<br>(0.917) |  |  |  | PTFE | 0.651<br>(0.651) |  |  |  |
| sandpaper | 0.579<br>(0.463) | 0.579<br>(0.461) |  |  | sandpaper | 0.012*<br>(0.010) | 0.012*<br>(0.011) |  |  |
| hook | 0.024*<br>(0.010) | 0.024*<br>(0.010) | 0.030*<br>(0.018) |  | hook | 0.011*<br>(0.007) | 0.011*<br>(0.007) | 0.011*<br>(0.007) |  |
| loop | 0.024*<br>(0.010) | 0.024*<br>(0.010) | 0.030*<br>(0.018) | 1.000<br>(1.000) | loop | 0.011*<br>(0.007) | 0.011*<br>(0.007) | 0.011*<br>(0.007) | NA |

For 10-DO larvae, the hook and loop treatments showed 2 ± 2% (mean ± standard error (SE)) and 3 ± 3% escape, respectively, which were not statistically different from each other (Figure 3A; Table 3). These two treatments were significantly more effective than the untreated (51 ± 12.4%), PTFE (36 ± 9.0%), and sandpaper (34 ± 10.7%). The escape rates of the untreated, PTFE, and sandpaper were comparable to each other.

For 20-DO larvae, both the hook and loop treatments resulted in 0 ± 0% escape, again significantly lower than those of the others (untreated: 94 ± 2.45%, PTFE: 92 ± 2.55%, sandpaper: 78 ± 1.22%; Figure 3B; Table 3). Unlike for 10-DO larvae, the sandpaper treatment was also more effective than the untreated and PTFE, while the untreated and PTFE did not differ from each other.

In Experiment 2, custom-designed anti-climbing tapes with protrusions effectively prevented larval escape, although the effect diminished as the protrusion gap sizes increased (Figure 4; see Table 4 for *P*-values).

**FIGURE 4.**
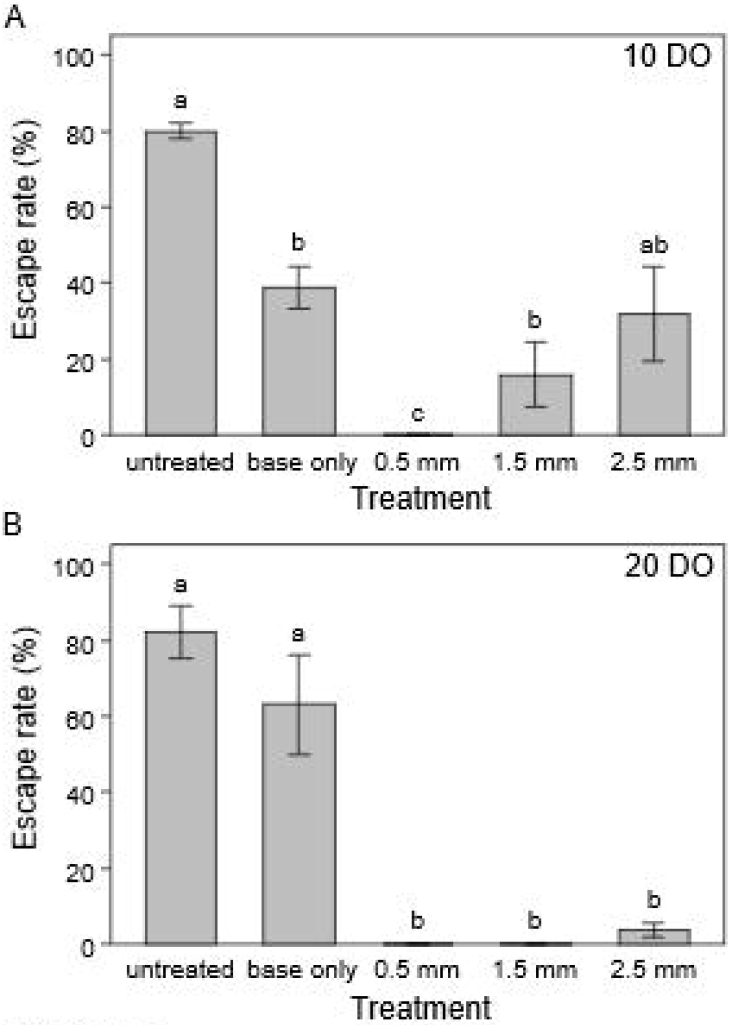
Escape rates of BSFL from containers treated with various protrusion gap sizes. (A) 10-DO larvae. (B) 20-DO larvae. Data were presented as mean ± standard error (*n* = 5). 20 individuals were used per replication. Different letters above bars indicate significant differences (*P* < 0.05) according to pairwise Wilcoxon rank sum test with Benjamini-Hochberg correction.

**TABLE 4.** Summary of multiple comparisons between larval escape rates from rearing containers treated with anti-climbing tapes of different protrusion gap sizes. *P*-values derived from pairwise Wilcoxon rank sum test with Benjamini-Hochberg correction are shown. Values inside parentheses indicate *P*-values without correction *: *P*< 0.05, NA: test not available (complete coincidence of values between groups).

| 10 DO | untreated | base only | 0.5 mm | 1.5 mm | 20 DO | untreated | base only | 0.5 mm | 1.5 mm |
| --- | --- | --- | --- | --- | --- | --- | --- | --- | --- |
| base only | 0.023*<br>(0.011) |  |  |  | base only | 0.249<br>(0.249) |  |  |  |
| 0.5 mm | 0.023*<br>(0.007) | 0.023*<br>(0.007) |  |  | 0.5 mm | 0.017*<br>(0.007) | 0.017*<br>(0.007) |  |  |
| 1.5 mm | 0.023*<br>(0.011) | 0.070<br>(0.056) | 0.041*<br>(0.025) |  | 1.5 mm | 0.017*<br>(0.007) | 0.017*<br>(0.007) | NA |  |
| 2.5 mm | 0.064<br>(0.045) | 0.260<br>(0.234) | 0.023*<br>(0.007) | 0.293<br>(0.293) | 2.5 mm | 0.018*<br>(0.012) | 0.018*<br>(0.012) | 0.080<br>(0.071) | 0.080<br>(0.071) |

For 10-DO larvae, the 0.5-mm gap treatment reduced escape to 0 ± 0%, greatly outperforming both the untreated (80 ± 2.24%) and base-only (39 ± 5.57%) controls (Figure 4A; Table 4). The escape rate increased with gap size to 16 ± 8.43% (1.5-mm; marginally significant) and 32 ± 12.3% (2.5-mm). The 1.5-mm gap was more effective than the untreated but not the base-only, while the 2.5-mm gap was not more effective than either. The base-only, however, did reduce escape compared to the untreated.

For 20-DO larvae, all protrusion treatments were highly effective (Figure 4B; Table 4). The 0.5-mm and 1.5-mm gaps both resulted in 0 ± 0% escape. Although the 2.5-mm gap showed 4 ± 1.87% escape, this did not differ from the other gaps. All three rates were significantly lower than those of the untreated (82 ± 6.82%) and base-only (63 ± 13%) controls. Unlike for 10-DO larvae, the base-only treatment was statistically indistinguishable from the untreated.

In Experiment 3, anti-climbing tapes with a greater number of protrusion rows tended to be more effective at preventing larval escape, when the gap size was held constant (Figure 5; see Table 5 for *P*-values).

**FIGURE 5.**
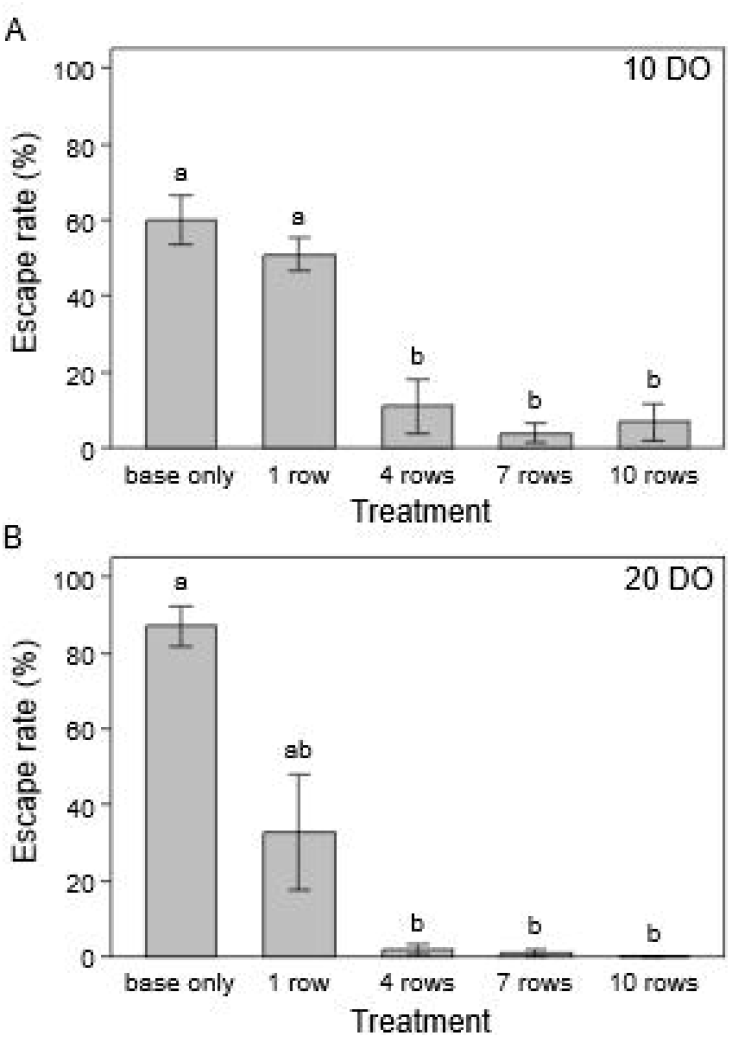
Escape rates of BSFL from containers treated with various protrusion row numbers. (A) 10-DO larvae. (B) 20-DO larvae. Data were presented as mean ± standard error (*n* = 5). 20 individuals were used per replication. Different letters above bars indicate significant differences (*P*< 0.05) according to pairwise Wilcoxon rank sum test with Benjamini-Hochberg correction

**TABLE 5.** Summary of multiple comparisons between larval escape rates from rearing containers treated with anti-climbing tapes of different protrusion row numbers. *P*-values derived from pairwise Wilcoxon rank sum test with Benjamini-Hochberg correction are shown Values inside parentheses indicate *P*-values without correction *: *P*< 0 05.

| 10 DO | base only | 1 row | 4 rows | 7 rows | 20 DO | base only | 1 row | 4 rows | 7 rows |
| --- | --- | --- | --- | --- | --- | --- | --- | --- | --- |
| 1 row | 0.850<br>(0.595) |  |  |  | 1 row | 0.073<br>(0.037) |  |  |  |
| 4 rows | 0.022*<br>(0.011) | 0.024*<br>(0.014) |  |  | 4 rows | 0.036*<br>(0.011) | 0.145<br>(0.101) |  |  |
| 7 rows | 0.022*<br>(0.011) | 0.022*<br>(0.011) | 0.904<br>(0.723) |  | 7 rows | 0.036*<br>(0.010) | 0.096<br>(0.057) | 0.600<br>(0.600) |  |
| 10 rows | 0.022*<br>(0.011) | 0.022*<br>(0.011) | 0.906<br>(0.906) | 0.906<br>(0.905) | 10 rows | 0.036*<br>(0.007) | 0.063<br>(0.025) | 0.221<br>(0.177) | 0.471<br>(0.424) |

For 10-DO larvae, the 1-row treatment resulted in 42 ± 7.68% escape, which was not significantly different from the base-only control (60 ± 6.33%; Figure 5A; Table 5). In contrast, treatments with 4, 7, and 10 rows reduced escape to 11 ± 7.14%, 4 ± 2.45%, and 7 ± 4.90%, respectively. These multi-row treatments were significantly more effective than both the base-only and the 1-row treatment. No significant differences were detected among the 4-, 7-, and 10-row treatments.

For 20-DO larvae, the 1-row treatment again showed an escape rate (33 ± 15.2%) that was not significantly different from the base-only control (87 ± 5.39%; Figure 5B; Table 5). The 4-, 7-, and 10-row treatments highly reduced escape to 2 ± 1.23%, 1 ± 1%, and 0 ± 0%, respectively. All three rates were significantly lower than that of base-only. Unlike for 10-DO larvae, however, these multi-row treatments were not statistically different from the 1-row treatments. Similar to the 10-DO group, no significant differences were confirmed among 4-, 7-, and 10-row treatments.

In Experiment 4, by applying anti-climbing tapes to rearing containers, we observed zero larval escape throughout the 20-day rearing period using 10-DO larvae (Figure 6). In contrast, the untreated containers displayed a final cumulative escape rate of 53.79 ± 3.77% (349.64 ± 24.5 out of 650 individuals). From the untreated containers, larval escape occurred in two distinct phases. First, the escape rate reached 32.31 ± 1.03% (210 ± 6.68 individuals) by day 1 and then remained relatively stable until day 14 with less than five escapes per container per day. Then from day 15, coinciding with the first prepupal escape, the rate increased sharply until the end of trials, averaging 21 escapes per container per day.

**FIGURE 6.**
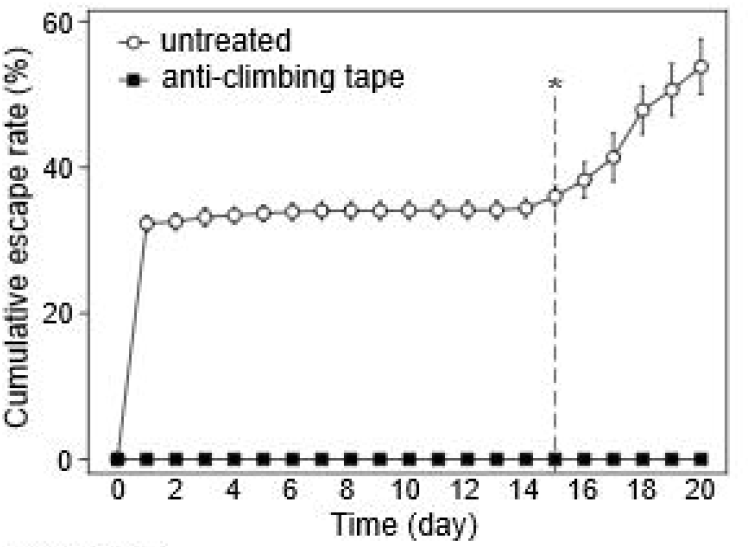
Cumulative escape rates of BSFL in a large-scale rearing setup. The anti-climbing tapes consisted of 15-row protrusions with 0.5-mm gaps. Data were presented as mean ± standard error (*n* = 3). 650 individuals (10-days-old) were used per replication. No statistical analysis was conducted. An asterisk indicates the day when the first prepupal escape was observed (Day 15).

## Discussion and conclusion

To travel on vertical walls, apodous BSF larvae rely on adhesion to the substrate. On smooth surfaces of rearing containers typically made of plastics such as polypropylene, this is likely achieved through wet adhesion arising from a thin liquid film or bridge between the larval cuticle and the wall surface (Supplementary Figure S3A; Butt & Kappl, 2009; Dirks, 2014). This wet adhesion generates an adhesive force that pulls the larval body toward the wall surface. The resulting normal force between the larva and the wall is a prerequisite for generating the frictional forces necessary for upward propulsion. Therefore, adhesive force likely plays a central mechanical role in enabling vertical climbing. Because the capillary force — the major component of wet adhesive force — is widely considered to be approximately proportional to the liquid-mediated contact area (Butt & Kappl, 2009; Langowski *et al*., 2018), we initially hypothesised that reducing the physical contact area between the larvae and the container wall would sufficiently weaken adhesive force to prevent larval climbing and escape.

To test this hypothesis, we first compared larval adhesive force across several materials and found that surfaces with protrusions (sandpaper and hook-and-loop fasteners) substantially reduced the adhesive force, even relative to the low-wettability surface of PTFE (Dhanumalayan and Joshi, 2018; Figure 2A). However, given that only the hook-and-loop fasteners exhibited a clear escape-prevention effect in the subsequent assessment (Figure 3), these results cannot be solely explained by differences in adhesive force.

Therefore, we revised our initial hypothesis to propose that, in addition to weakened adhesion, the three-dimensional structure of macroscale protrusions (a few millimeters in height) may mechanically hinder larval climbing. Specifically, we postulated that an array of protrusions on the wall would require larvae to lift their posterior body to reach the top of protrusions. This postural change may increase gravitational torque promoting backward flipping (Günther and Weihmann, 2012; Ritzmann *et al*., 2005), while simultaneously decreasing the counteracting torque due to weakened wet adhesion on the anterior body, thereby rendering larval climbing kinematically unstable.

Based on this idea, we devised anti-climbing tapes — novel tape-like structures with regularly arranged macroscale protrusions on their surfaces (Figure 1; Table 1). The protrusions were designed in a truncated-cone shape to improve durability and facilitate cleaning, both of which are limited in conventional hook-and-loop fasteners. Surfaces with protrusions significantly reduced larval adhesive force compared to flat surfaces without protrusions (Figure 2B), likely due to a reduction in effective contact area.

Further investigation into the design parameters provided evidence that this method operates through a dual mechanism: first, by imposing a direct physical constraint on larval movement, and second, by reducing adhesion, as initially hypothesised.

The first mechanism, physical constraint, becomes critical when larvae attempt to enter the gaps between protrusions. Larvae are blocked at the lower edge of the tapes when the gaps are too narrow for them to squeeze through (Supplementary Figure S3B, E), whereas they could readily pass through when the gaps are wide enough, often also gaining mechanical support (Supplementary Figure S3C, D). Indeed, within approximately 1 hour after larvae were introduced into containers treated with the tapes, the majority of larvae attempting to escape did not climb over the protrusions but instead tried to squeeze into the gaps between them. However, by 72 hours after the beginning of the experiment, substantial escape was observed (Figure 4; comparison between ‘base only’ and ‘1 row’). These findings suggest that this mechanism alone is insufficient to ensure long-term escape prevention.

The second mechanism, adhesion reduction, becomes critical when larvae attempt to climb over the protrusions. When larvae fail to enter the gaps, they extend their bodies across the protrusion tops (Supplementary Figure S3F, G); however, the reduced contact area on these elevated surfaces decreases effective adhesion. At the same time, because the protrusion tops are elevated relative to the base surface, the distance between the center of mass and the wall increases, thereby increasing the gravitational torque described above. Importantly, the effectiveness of this process likely depends on the width of the protrusion-top region, which is determined by the number of protrusion rows. If this region is shorter than the larval body length, the anterior end may reach the adjacent flat surface before torque-induced destabilisation becomes sufficient to cause backward rotation, partially restoring whole-body adhesion and reducing the risk of detachment. In this way, the escape-prevention efficacy is determined by both adhesion reduction and the vertical extent of the protrusion array.

These findings suggest that a sufficiently wide protrusion array is required to force the entire body of the larva onto the low-adhesion surface. Indeed, on wider arrays (*e.g*., 10-row tapes), the adhesion-reduction mechanism was effective for larger 20-DO larvae, which were unable to escape (Figure 5). However, some smaller 10-DO larvae still managed to traverse the array and escape, suggesting they were able to generate sufficient adhesion on the limited top surfaces. This size-dependent escape pattern indicates that smaller larvae may be less affected by the reduction in contact area. This could be attributed to factors such as the more deformable cuticle of younger instars (Rebora *et al*., 2023) and higher surface-area-to-volume ratio, which may allow better conformational matching to the substrate and more efficient adhesion per unit body mass (Labonte and Federle, 2015; Langowski *et al*., 2018). Although further reducing the top surface area could potentially mitigate this issue, technical constraints of our fabrication system limited the minimum diameter to 0.9 mm in the present study.

Finally, we validated a design featuring 0.5-mm gaps and 15 rows in a large-scale, 20-day rearing trial. In all three replicates, this design completely prevented larval escape from the fourth to fifth instar (approximately 10 mm in length) through the prepupal stage (Figure 6), confirming its practical applicability for rearing later-instar larvae. To better define the operational limits of the method, however, we conducted additional exploratory trials that were not part of the formal experimental design. Using a larval group that included smaller individuals (2–10 mm in length), we found that the 15-row protrusions did not fully prevent escape. As inferred above, this outcome is likely attributable to their smaller body size, which may confer two advantages: reduced sensitivity to contact-area limitation and enhanced ability to mechanically interlock within the protrusion gaps (Supplementary Figure S3H).

In conclusion, we demonstrate that an array of protrusions prevents BSF larval escape via a dual mechanism: physical obstruction blocks larvae from passing through the gaps, while adhesion reduction hinders them from climbing over the array. To maximise efficacy, design parameters must address both aspects: (1) gaps must be narrow enough to physically block the target larvae; (2) the protrusion array must extend over a sufficient vertical span to prevent larvae from traversing it; and (3) the top surface area of each protrusion should be minimised (*e.g*., ≤ 0.9 mm in diameter) to further limit adhesion. Furthermore, we recommend applying this method primarily to larvae larger than 10 mm in length to ensure reliable performance.

