## Supplementary Figures for "A novel anti-climbing barrier prevents black soldier fly larval escape from rearing containers"

RESEARCH ARTICLE

**Supplementary material**

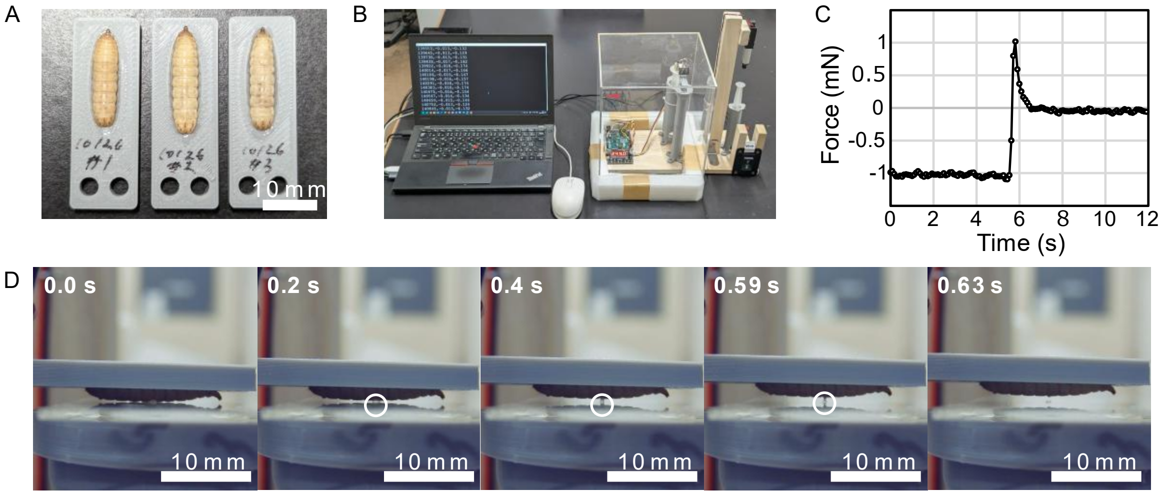


FIGURE S1. Overview of the adhesive force measurement using black soldier fly larvae. (A) Larvae fixed onto plates for measurement. (B) Overall view of the measurement device. (C) An example of a measurement result (glass substrate). (D) Deformation of the water capillary bridge observed during measurement. As the larva is pulled away from the substrate, the bridge gradually elongates and eventually ruptures.

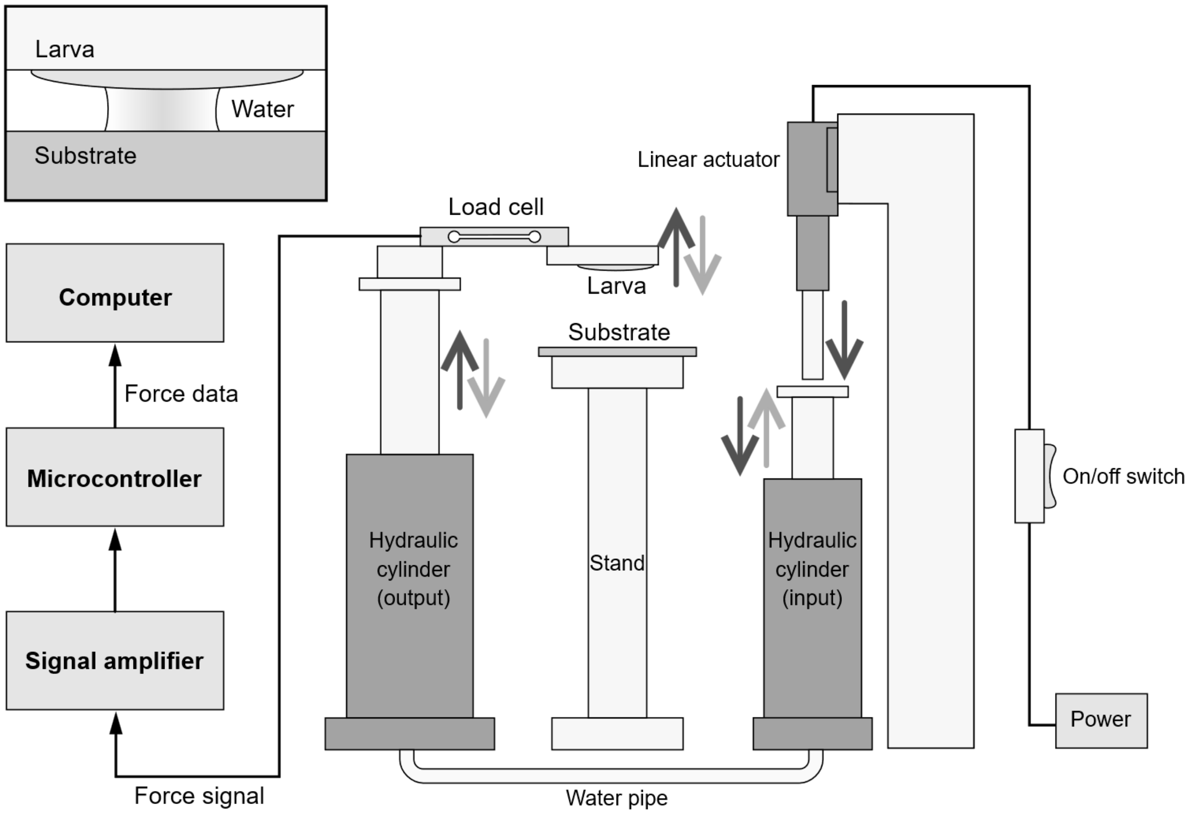


FIGURE S2. Schematic overview of the adhesive force measurement device used in this study. Open arrows indicate the direction of movement of each movable component; arrows of the same color move synchronously. The inset in the upper left shows a schematic representation of the capillary bridge formed between the larva and the substrate during measurement. The sizes of the components in the figure are not drawn to scale.

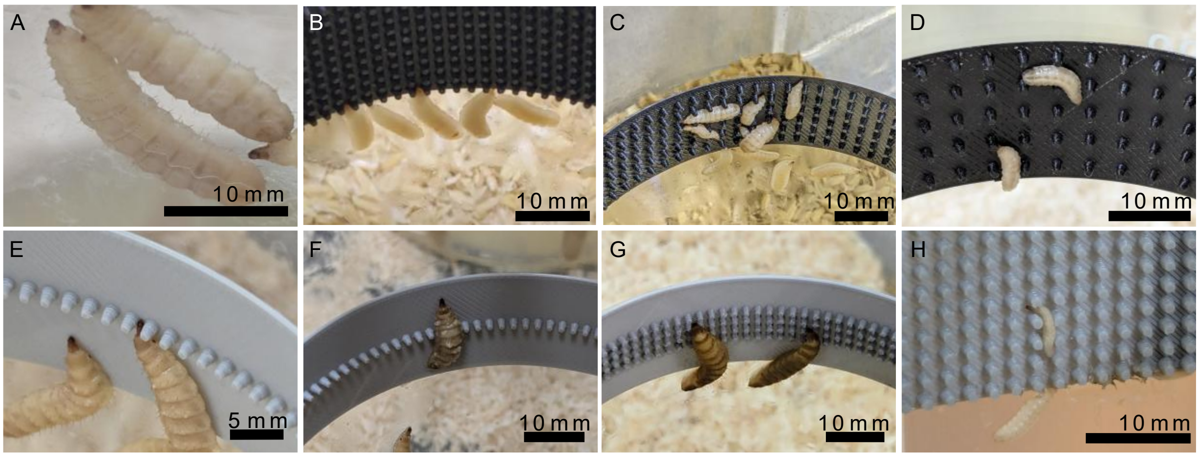


FIGURE S3. Photographs of black soldier fly larvae on vertical surfaces. (A) Thin film of water formed between a glass wall and larval cuticle. Larvae responding to anti-climbing tapes with (B) 0.5-mm, (C) 1.5-mm, and (D) 2.5-mm gap sizes. (E) A larva trying to pass through a gap between protrusions (gap size: 0.5 mm, row number: one). (F) A larva traversing the row of protrusions (gap size: 0.5 mm, row number: 1). (G) Larvae trying to climb over the array of protrusions (gap size: 0.5 mm, row number: 4). (H) A small larva (<< 10 mm in length) climbing through gaps between protrusions in an additional explanatory trial (gap size: 0.5 mm, row number: 15).
